# Expression of IL-1R2 by T follicular regulatory cells prevents the exacerbation of allergy by blocking their IL-1-dependent proliferation

**DOI:** 10.1101/2024.08.29.610028

**Authors:** Paul Engeroff, Aude Belbezier, Romain Vaineau, Gwladys Fourcade, Hugo D Lujan, Bertrand Bellier, Stephanie Graff-Dubois, David Klatzmann

## Abstract

The antibody response is regulated by follicular T helper (Tfh) and regulatory (Tfr) cells that control the germinal center (GC) reaction. Recent research has shown that Tfh/Tfr have a unique pattern of IL-1 receptor expression. We investigated the mechanisms by which this IL-1 axis in GCs could regulate the allergic response.

To study this, we generated CD4^cre^IL-1R1^lox^ mice, specifically lacking IL-1R1 expression in T cells and FoxP3^cre^IL-1R2^lox^ mice, specifically lacking IL-1R2 expression in Tfr cells. The conditional knockout mice were compared to their respective control mice in a model of ovalbumin (OVA) sensitization and anaphylaxis, and a phenotypic and functional characterization of humoral and cellular responses was performed.

While CD4^cre^IL-1R1^lox^ mice showed little phenotypic changes, FoxP3^cre^IL-1R2^lox^ mice were highly susceptible to allergic anaphylaxis and generated an increase in IgE responses that promoted basophil degranulation. Additionally, FoxP3^cre^IL-1R2^lox^ mice displayed significantly reduced OVA-specific IgG responses, limiting their ability to control allergy via the inhibitory IgG receptor FcγRIIb. Although FoxP3^cre^IL-1R2^lox^ mice showed an overall increase in splenic T and B cell numbers, they were unable to efficiently generate proliferating GC B cells. Upon *ex vivo* IL-1β and/or OVA re-stimulation, we observed a striking IL-1R1-dependent activation and proliferation of Tfr cells in FoxP3^cre^IL-1R2^lox^ splenocytes, that was neither observed in Tregs nor in Tfh. At the same time, B cell proliferation upon re-stimulation was suppressed.

These findings suggest that IL-1R2 expression on Tfr cells prevents allergy by limiting excessive Tfr activation and suppressing the IgG/IgE ratio.

## INTRODUCTION

Type I hypersensitivities including allergic airway diseases, food allergy and severe cases of allergic anaphylaxis are on the rise worldwide^1,2^. Type I hypersensitivity is characterized by a misguided immune response resulting in elevated levels of Immunoglobulin E (IgE). IgE sensitizes mast cells and basophils via the high-affinity IgE receptor FcɛRI, which gears the cells up to de-granulate upon allergen encounter resulting in the release of highly inflammatory mediators^2–6^.

In recent years, it has become clear that IgG antibodies play a critical role in regulating the inflammatory activity of IgE. Allergen-specific IgG can either neutralize the allergen, or inhibit FcɛRI signalling directly on mast cells/basophils by engaging the IgG receptor FcγRIIb^7–12^. A protective role for IgG against allergy has been observed in natural allergen tolerance and acquired allergen tolerance through allergen-specific immunotherapy (AIT)^13–17^. Monoclonal IgG anti-allergen antibodies represent a promising therapeutic strategy for the same reason^18–20^. Thus, the balance between IgG and IgE is an important regulator of allergic sensitization.

However, the mechanisms that control the IgG versus IgE response to allergen immunization are poorly understood. Allergen-specific antibody responses are controlled by T follicular helper cells (Tfh) and T follicular regulatory cells (Tfr) during the germinal center (GC) reaction, the process by which B cells undergo affinity maturation to generate high-affinity antibody responses^21^. Even though the specific Tfh/Tfr-mediated mechanisms that shape the allergic response are not yet fully resolved, it is becoming clear that they are critical regulators of allergic sensitization^22–28^.

Our group has recently described an IL-1 axis that controls the activation of Tfh and Tfr cells during the GC reaction, which is driven by their unique IL-1 receptor (IL-1R) expression pattern^29–31^. Tfh cells express the IL-1R1 agonist receptor, whereas Tfr cells express lower levels of IL-1R1, but high levels of the inhibitory IL-1R2 decoy receptor^29–31^. How these IL-1Rs in Tfh/Tfr shape the IgG/IgE ratio and allergic sensitization is poorly understood.

Here, we identify that the deletion of IL-1R2 expression in Tfr cells activates their own IL-1R1-dependent proliferation, leading to a disruption of GC B cell proliferation, which creates an allergic phenotype in mice characterized by a suppression of the IgG/IgE ratio.

## MATERIAL AND METHODS

### Mice

All animals were kept at the Centre d’Expérimentation Fonctionnelle animal facility (Paris, France). The local animal ethics committee approved all procedures. Transgenic mouse strains CD4^cre^IL-1R1^lox^ and FoxP3^cre^IL-1R2^lox^, and their respective control lines as well as their basic naïve phenotypes were previously described^30^. The experiments shown were done with 6–8-week-old female mice. For some repeat experiments (not shown) female mice were used for experiments up to 15 weeks old. The groups within individual experiments were performed with female mice at the same age (± 7 days).

### Injections

All injections except OVA challenges were done in a volume of 100ml. Mice were immunized day 0 and day 14 by subcutaneous injection with 10µg OVA (OVA A5503, Sigma-Aldrich) in Imject® Alum (Thermo Fisher) 1:1 diluted in PBS. For induction of anaphylaxis, mice were injected intravenously with 2 μg OVA in 200ml PBS. Rectal temperature was measured at 10-minute intervals for 1 hour using a rectal thermometer (Biosep Lab Instruments, Vitrolles, France). For blocking of FcγRIIb, 100μg anti-CD32b antibody (Thermo Fisher, clone AT130-2) was injected intraperitoneally 1 hour before OVA challenge. Alternatively, Mouse IgG2a, kappa Isotype control (Miltenyi Biotec) was injected.

### Sampling

To assess basophils, blood from tail veins was collected in EDTA tubes (final concentration 1mM EDTA). Red blood cells were lysed using BD Pharm Lyse™ (BD Bioscience) according to the manufacturers protocol and were washed three times with PBS. Whole spleens were collected from sacrificed mice and transferred into 48-well plates containing RPMI 1640 medium (Sigma). The spleens were then mashed and filtered through a cell strainer (70μm, Sigma Aldrich) and washed three times with PBS and were finally suspended in RPMI 1640, 20% FCS medium (Sigma).

### Flow cytometry

All primary antibody stainings were carried out in 96 round well plates (Thermo Fisher) for 20 minutes at 4°C in PBS. Between all steps, the cells were washed two times with 200µl PBS. Fixation and permeabilization was performed with eBioscience™ Intracellular Fixation & Permeabilization Buffer Set (Thermo Fisher) according to the manufacturers protocol. Basophils were defined as anti-mouse CD49b+ (Thermo Fisher, clone DX5), anti-mouse IgE+ (Thermo Fisher, clone 23G3), anti-mouse CD45lo (Thermo Fisher, clone 30-F11) and negative for anti-CD117 (Thermo Fisher, clone 2B8) and anti-CD19 (BD Bioscience, clone 1D3). Basophil activation was investigated with anti-CD63 (Thermo Fisher, clone NVG-2) staining. Germinal center B cells were defined as CD45+, anti-mouse CD19+ or anti-mouse B220+ (Thermo Fisher, clone RA3-6B2), anti-mouse GL-7+ (Thermo Fisher, clone GL-7) anti-mouse IgD-(Thermo Fisher, clone 11-26c) and anti-mouse CD3-(Thermo Fisher, clone 145-2C11). OVA-specific GC B cells were investigated by staining with 1µg/ml Invitrogen™ Ovalbumin, Alexa Fluor™ 488 Conjugate (Thermo Fisher Scientific). Proliferating cells were investigated by staining with anti-mouse Ki-67 (Thermo Fisher, clone SolA15). Plasmablast were defined as CD19+/B220+, GL-7-, IgD-, CD138+ (Thermo Fisher, clone 281-2). Follicular T cells were gated by anti-mouse CD3+, anti-mouse CD4+ (Thermo Fisher, clone RM4-5) anti-mouse CXCR5+ (Thermo Fisher, clone 2G8) and anti-mouse PD-1+ (Thermo Fisher, clone RPM1-30) dumping CD19+ or B220+ cells. In Follicular T cells, Tfh cells were defined as anti-mouse FoxP3-(Thermo Fisher, clone FJK-16s) whereas Tfr were defined as FoxP3+, anti-mouse CD69 (BD Bioscience, clone H12F3), and anti-mouse ICOS (Thermo Fisher, clone 7E.17G9). All flow cytometry was performed with CytoFLEX (Beckman Coulter, Brea, USA) and analyzed by using FLOWJO software (TreeStar Inc, Ashland, Ore) or CytEXPERT (Beckman Coulter).

### ELISA

For ELISA, 96-well Nunc Maxisorp ELISA plates (Thermo Fisher Scientific) were coated with reagents in PBS at 4°C overnight. For anti-OVA IgG, 500ng/ml OVA was coated. For total and specific IgE, rat anti-mouse anti-IgE (BD Biosciences, clone R35-72) was coated at 2µg/ml. Blocking was performed with Blocker^TM^ Casein solution (Thermo Fisher) for 2 hours. Dilutions of sera were added to the plates and incubated for 2 hours at room temperature. For specific IgG ELISA, serum was initially diluted 1:100, followed by 10x dilution steps. For total IgE, initial dilution was 1:50 followed by 3x dilution steps. To generate a standard curve, we used purified mouse IgE (Biolegend, clone MEA-36). For specific IgE, serum dilution was 1:10. The plates were then washed 5 times with PBS/0.05% Tween. Specific IgG was detected for 1 hour at room temperature with biotin rat anti-mouse IgG (Southern Biotech, Birmingham, AL, USA) biotin rat anti-mouse IgG1 (Southern Biotech), or biotin rat anti-mouse IgG2b (Southern Biotech). Total IgE was detected with biotin rat anti mouse IgE (Southern Biotech). After washing 3 times in PBS/0.05% Tween, specific IgG and total IgE ELISAs were further incubated with Streptavidin-HRP (Thermo Fisher) for 1 hour at room temperature. Specific IgE was detected with 500ng/ml OVA added for 1 hour at room temperature in a first step. Plates were washed 5 times with PBS-Tween and thereafter incubated with polyclonal anti-OVA HRP (Thermo Fisher). Finally all plates were washed 5 times with PBS/0.05% Tween and the ELISAs were developed with 1x TMB substrate solution (Thermo Fisher) and stopped with 1M HCL. All ODs were measured at 450 nm, the half-maximal antibody titer (OD_50_) was defined as the reciprocal of the dilution leading to half of the OD measured at saturation. To visualize the changes in antibody composition, we estimated an IgG/IgE ratio. Due to the low levels of detectable specific IgE in the serum, we generated a ratio between OD_50_ IgG titer and OD_450_ IgE signal, measured from the same serum samples. For avidity assays, an additional washing step in PBS/0.05% Tween or 7 M urea, three times for 5 min at room temperature was included after incubation with serum. The avidity index is the urea/no-urea OD450 ratio.

### Effector cell binding and activation assays

For basophil assays, about 200ul blood from three mice was pooled and lysed and cells were resuspended in RPMI1640 medium with 10%FBS (Sigma). For basophil activation tests, cells were initially incubated with OVA for 1 hour at 37°C in. From titration experiments, 5nM OVA was established as a good dose for comparing basophil activation and used for all proceeding experiments. However, the basophil assays were later optimized by shortening the incubation time to 20 minutes at 37°C. The presence of mouse BD Fc-Block^TM^ (BD Biosciences) further allowed us to determine the IgG-dependent binding effect from our sera. For basophil activation tests, serum from D21 immunized mice was pre-mixed for 20 minutes at room temperature with 5nM OVA prior to addition to basophils. Binding assays in basophils were performed by premixing 25nM OVA-A488 with 1:10 diluted serum in medium for 20 minutes at 4°C. Activation/Inhibition assays were performed with 1:1000 diluted serum after initial titration experiments. For all activation assays, instead of Fcγ-block, the more specific blocking antibody anti-FcγRIIb (Thermo Fisher, clone AT130-2) was added at 1:200 dilutions to the cells prior to the addition of OVA-IgG complexes.

### T cell re-stimulation assay

Tfh/Tfr cells, re-stimulation experiments were performed with whole splenocytes derived from mice immunized with OVA at day 0 and 14. The spleens were collected at day 21 from either wild type mice or FoxP3^cre^IL-1R2^lox^ mice and splenocytes were isolated. The cells were cultured at a concentration of 2 million cells/mL in 96 well round (U) bottom plates (Thermo Fisher Scientific). Culture medium was RPMI 1640 (Thermo Fisher Scientific) supplemented with 10% FBS, 20mM L-Glutamine (Thermo Fisher Scientific) and Penicillin-streptomycin (100 U/mL, Thermo Fisher Scientific) in a final volume of 200µl per well. We based the IL-1 concentrations on our previous *in vitro* experiments where we used IL-1β to stimulate T cells^31^. Additionally, we aimed to use approximately equimolar ratios of IL-1β, OVA and Anakinra. Together, this led us to use 0.5 μg/ml IL-1β, 1.2 μg/ml OVA and 0.5 μg/ml Anakinra for the re-stimulation assays. The splenocytes were cultured for 48h at 37°C, 5% CO_2_ and the cells were then washed 3 times in PBS prior to staining.

### Statistics

All results shown are representative of at least 3 independent experiments. All data is presented as mean plus/minus the standard error of the mean. All the statistical tests and evaluations were performed by using GraphPad PRISM 6.0 (GraphPad Software, Inc, La Jolla, Calif). For all experiments, we used an α value of 0.05 and statistical significance for p-values is displayed as follows: ∗ is less than or equal to .05; ∗∗ is less than or equal to .01; ∗∗∗ is less than or equal to .001. Non-parametric, 2-tailed Student t tests were used for the comparison of two groups (e.g. antibody titers, cell frequencies). Multiple comparisons (e.g. anaphylaxis, proliferation assays) were performed by 2-way ANOVA followed by Bonferroni correction to obtain individual p-values.

## RESULTS

### Sensitized CD4^cre^IL-1R1^lox^ mice do not display altered IgG/IgE ratios or anaphylactic responses

Within T cells, the IL-1 agonist receptor IL-1R1 is mainly expressed in T follicular cells, and within those generally more on Tfh than on Tfr. To study the role of IL-1R1 in T follicular cells, we generated a mouse line in which IL-1R1 is selectively knocked out in CD4+ T cells (CD4^cre^IL-1R1^lox^, Figure 1A). Their basic characteristics including naïve cell frequencies, and IL-1 receptor expression profile have recently been shown^30^.

**Figure 1:**
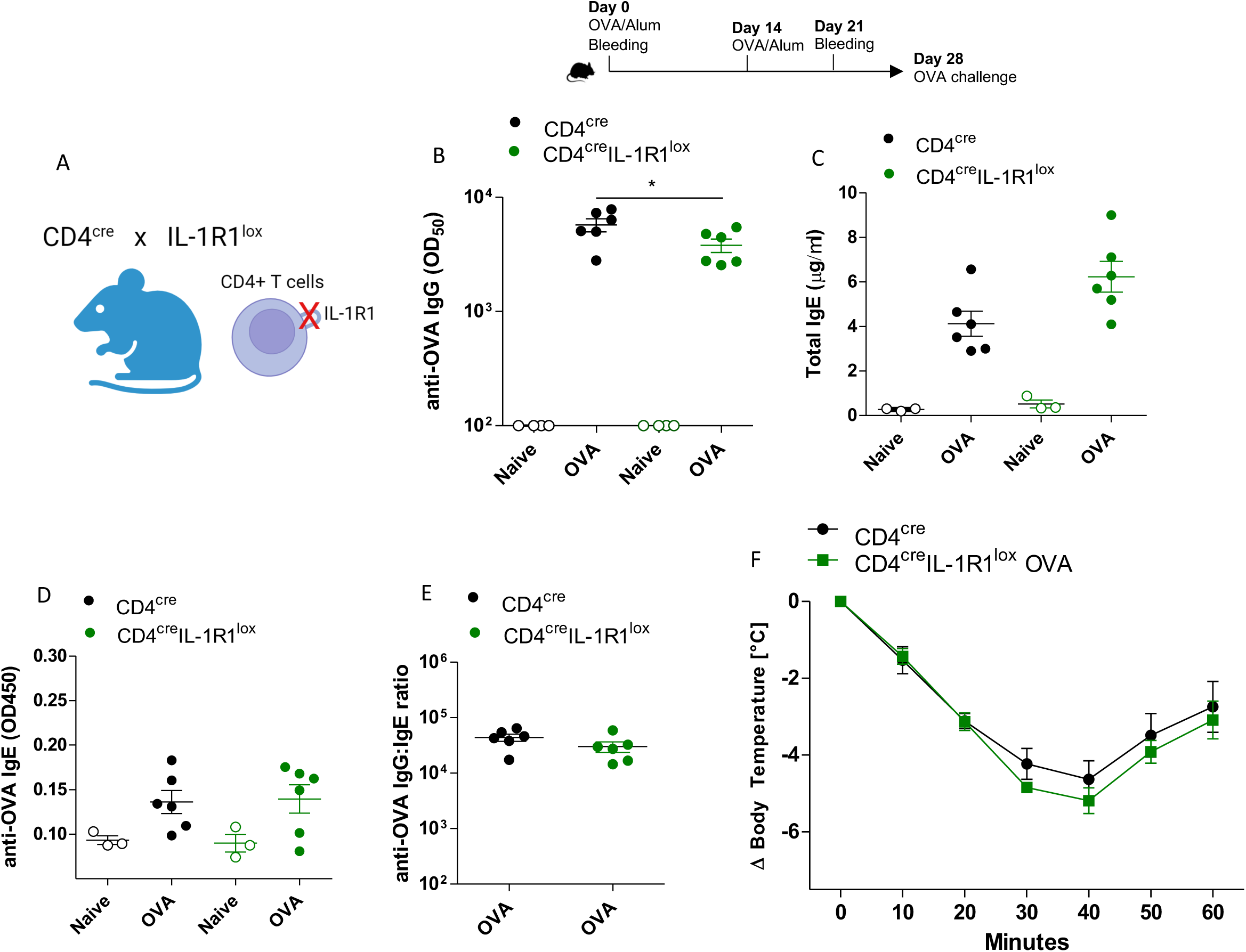
Sensitized CD4^cre^IL-1R1^lox^ mice do not display altered IgG/IgE ratios or anaphylactic responses. A) CD4^cre^IL-1R1^lox^ mice were generated. Mice were sensitized to OVA, serum was collected at day 0 and day 21, antibodies were determined by ELISA, OVA challenge was done at day 28. B) Shown are mean ± SEM OD_50_ titers of OVA-specific IgG. C) Mean ± SEM total IgE levels (µg/mL). D) Mean ± SEM OD450 values of OVA-specific IgE. E) Specific IgG to specific IgE ratio for individual mice. F) Shown are mean ± SEM degrees lost compared to baseline temperature upon OVA challenge (n=6/group).

We then tested these mice in an allergic context using an established ovalbumin (OVA)/alum sensitization model. Thus, we sensitized control (CD4^cre^) mice or CD4^cre^IL-1R1^lox^ with OVA and investigated IgG or IgE responses and the systemic anaphylactic response to OVA challenge. As shown in Figure 1B, we observe a reduction of OVA-specific IgG responses in CD4^cre^IL-1R1^lox^. Nevertheless, the mice still generated a strong IgG response compared to naïve mice, suggesting that OVA sensitization does not depend on IL-1R1 expression in T follicular cells.

We next investigated total IgE and OVA-specific IgE levels in those mice, both of which were induced compared to naïve mice by OVA/alum sensitization (Figure 1C, D). While there was a slight trend towards increasing total IgE levels in CD4^cre^IL-1R1^lox^ compared to control mice, the allergen-specific IgE response was overall not significantly changed. In recent years it has become clear, that the ratio between allergen-specific IgG and IgE is an essential determinant of the allergic response, as IgG can inhibit IgE effector functions^7^. We noted that the IgG/IgE ratio for individual mice did not significantly alter between CD4^cre^ controls and CD4^cre^IL-1R1^lox^ mice (Figure 1E).

Finally, we performed an intravenous OVA challenge, to investigate systemic anaphylaxis in those mice as measured by a drop of body temperature. As shown in Figure 1F, both CD4^cre^ controls and CD4^cre^IL-1R1^lox^ showed an anaphylactic response but we did not observe a significant difference in their responsiveness.

These findings suggest that the deletion of IL-1R1 expression in all CD4+ T cells, which is expected to mainly target T follicular cells, does not alter the allergic response.

### Sensitized FoxP3^cre^IL-1R2^lox^ mice display reduced IgG/IgE ratios and strongly enhanced anaphylaxis

We previously reported that non-follicular FoxP3+ T regs do not express IL-1 receptors, whereas Tfr express IL-1R1 and IL-1R2^31^. To study the role of IL-1Rs in Tfr, we previously crossed FoxP3^cre^ mice with IL-1R1^lox^ or IL-1R2^lox^ mice respectively. However, as FoxP3^cre^IL-1R1^lox^ mice still express IL-1R2, this may block their IL-1R1-dependent effects^30^. For these reasons, we here focused on FoxP3^cre^IL-1R2^lox^ mice which were sensitized to OVA/Alum and investigated for IgE response and anaphylaxis (Figure 2A).

**Figure 2:**
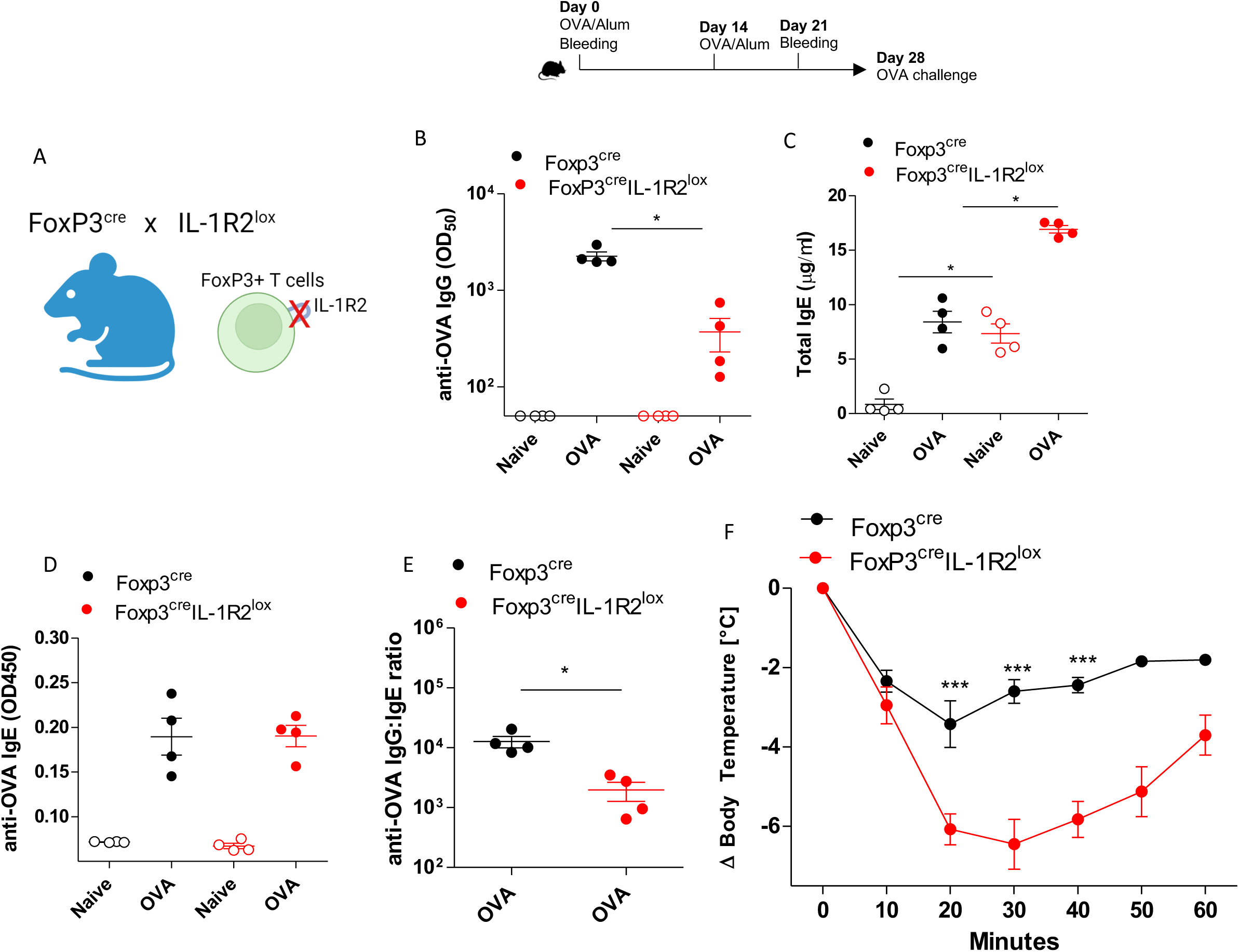
Sensitized FoxP3^cre^IL-1R2^lox^ mice display reduced IgG/IgE ratios and strongly enhanced anaphylaxis. A) FoxP3^cre^IL-1R2^lox^ mice were generated. Mice were sensitized to OVA, serum was collected at day 0 and day 21, antibodies were determined by ELISA, OVA challenge was done at day 28. B) Shown are mean ± SEM OD_50_ titers of OVA-specific IgG. C) Mean ± SEM total IgE levels (µg/mL). D) Mean ± SEM OD450 values of OVA-specific IgE. E) Specific IgG to specific IgE ratio for individual mice. F) Shown are mean ± SEM degrees lost compared to baseline temperature upon OVA challenge (n=4/group).

In these knockout mice, we observed a striking reduction of the OVA-specific IgG response compared to control mice (Figure 2B). Surprisingly, FoxP3^cre^IL-1R2^lox^ mice displayed significantly higher naïve total IgE levels that were even further increased upon sensitization (Figure 2C). As shown in Figure 2D, specific IgE levels were induced in both sensitized FoxP3^cre^ controls and in FoxP3^cre^IL-1R2^lox^ mice to equal levels. However, given the strong reduction of OVA-specific IgG, the specific IgG/IgE ratio for individual mice was overall significantly decreased (Figure 2E).

Finally, we challenged FoxP3^cre^IL-1R2^lox^ mice and their FoxP3^cre^ control mice to investigate systemic anaphylaxis. We noted a striking difference as FoxP3^cre^IL-1R2^lox^ were significantly more susceptible to systemic anaphylaxis upon allergen challenge, as evident from body temperature measurements (Figure 2F).

These findings suggest that the deletion of IL-1R2 expression in FoxP3+ cells, which is expected to specifically target Tfr cells, promotes an allergic phenotype characterized by a lower IgG/IgE balance and a significantly higher susceptibility to allergic anaphylaxis in response to allergen challenge.

### FoxP3^cre^IL-1R2^lox^ mice display increased basophil sensitization

While it is surprising that anaphylaxis in FoxP3^cre^IL-1R2^lox^ mice appears to occur independent of a change in allergen-specific IgE levels, past research has demonstrated that other features can regulate the allergic reaction. Higher IgE+ basophil frequencies and higher IgE densities on basophils can increase their sensitivity to degranulation^32,33^. To investigate the functional features of the IgE response in FoxP3^cre^IL-1R2^lox^ mice, we investigated basophil frequency, basophil IgE density and the activation of blood basophils by flow cytometry (Gating, Figure 3A).

**Figure 3:**
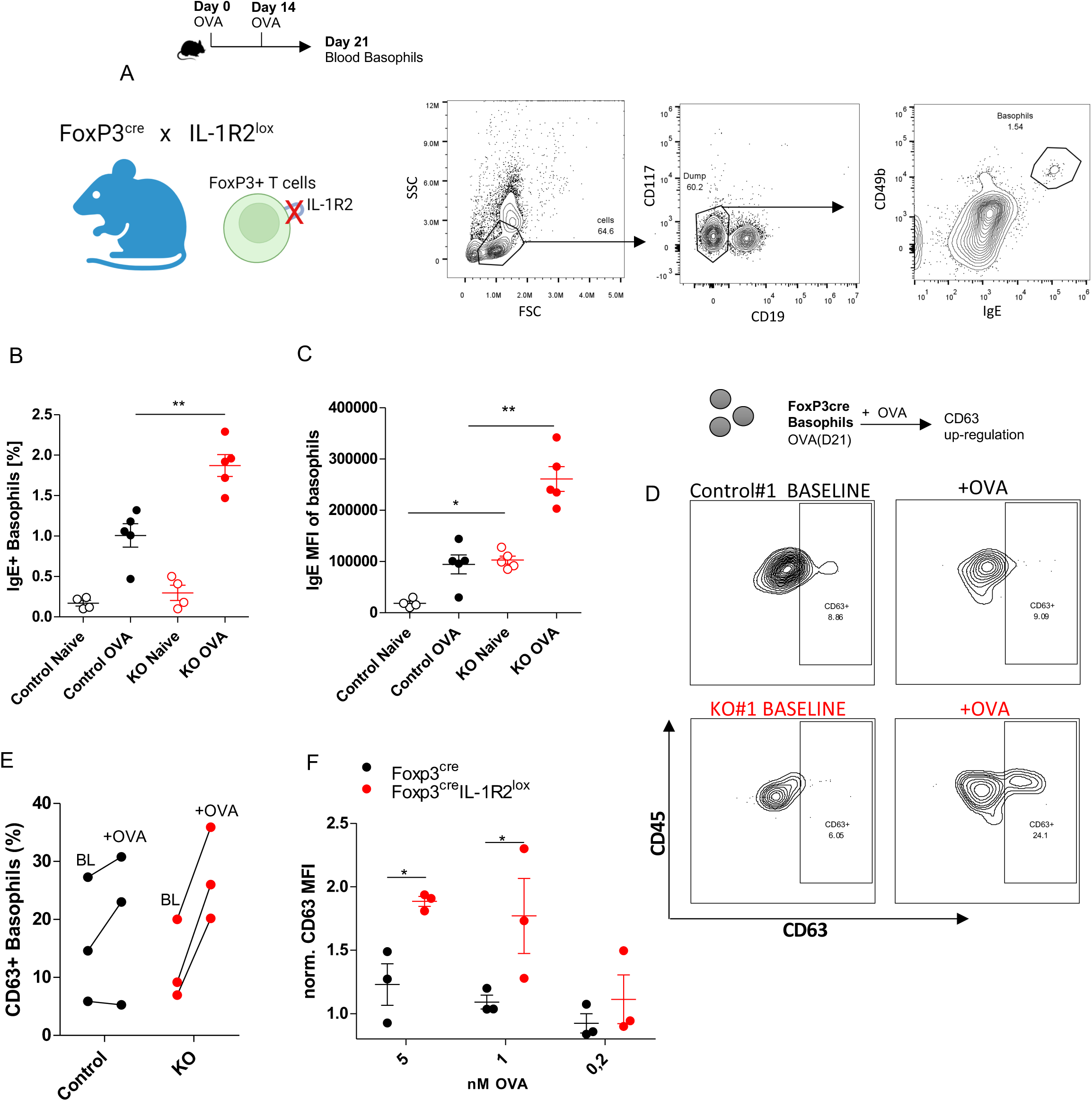
FoxP3^cre^IL-1R2^lox^ mice display increased basophil sensitization. FoxP3^cre^(Control) or FoxP3^cre^IL-1R2^lox^ (KO) mice were sensitized with OVA/Alum at day 0 and day 14, blood was collected at day 21. A) Shown is the basophil gating strategy B) Mean ± SEM % of IgE+ Basophils in CD45+ at day 21. C) Mean ± SEM basophil anti-IgE MFI at day 21. D-F) Basophils were challenged with OVA. D) Shown are basophil activation gating strategy and representative flow cytometry plots. E) Shown is he increase in % of CD63+ basophils from baseline (BL) values before challenge, to activated values (+OVA) after challenge. F) Shown are the mean ± SEM of anti-CD63 MFI values for titrated OVA doses normalized to respective baseline values.

The higher serum total IgE levels in FoxP3^cre^IL-1R2^lox^ mice translated to higher IgE+ blood basophil frequencies compared to control mice as well as increased surface IgE densities (Figure 3B, C). Comparative kinetics of total IgE and IgE+ basophils, as well as basophil surface IgG levels are shown in suppl. Figure 1. We next aimed to investigate if the increased basophil-displayed IgE corresponds to increased basophil degranulation upon OVA challenge.

We thus established basophil activation tests using OVA challenge for investigation by flow cytometry, staining for the degranulation marker CD63. Figure 3D shows raw flow cytometry plots of baseline CD63 staining compared to CD63 staining of OVA-challenged cells. In general, we noted a high variability in baseline CD63+ basophils, but as shown in Figure 3E, basophils from FoxP3^cre^IL-1R2^lox^ strongly up-regulated CD63 in response to OVA challenge whereas control mouse basophils were consistently less reactive. Normalization of the anti-CD63 MFI to baseline values resulted in a consistent readout for CD63 up-regulation (Figure 3F).

In conclusion, despite equal levels of serum OVA-specific IgE, the increased basophil-displayed IgE in FoxP3^cre^IL-1R2^lox^ mice favours basophil degranulation in response to OVA challenge. These results indicate that IL-1R2 expression by Tfr prevents systemic basophil sensitization in response to allergen immunization.

### FoxP3^cre^IL-1R2^lox^ mice fail to suppress systemic anaphylaxis and basophil degranulation via FcγRIIb

Having shown that overall basophil reactivity is higher in FoxP3^cre^IL-1R2^lox^ mice, we next aimed to investigate how the suppression of serum IgG can affect the allergic response. IgG is known to suppress IgE-dependent activation of allergic effector cells by various mechanisms^7^.

Thus, we next characterized the quality of anti-OVA IgG further by studying IgG subclasses and IgG avidity. The suppression of the IgG was global, as we noted a reduction across IgG1 and IgG2b subclasses and likewise a trend for a reduction of IgG avidity in FoxP3^cre^IL-1R2^lox^ compared to controls (Figure 4A-C). Of note, even though both were significantly lower, the magnitude of the observed reduction for IgG2b was more striking than for IgG1. These results imply that there is not only a shift in IgG/IgE ratios but that there is likewise a shift in IgG quality, as IgG subclasses and avidity are changed in FoxP3^cre^IL-1R2^lox^.

**Figure 4:**
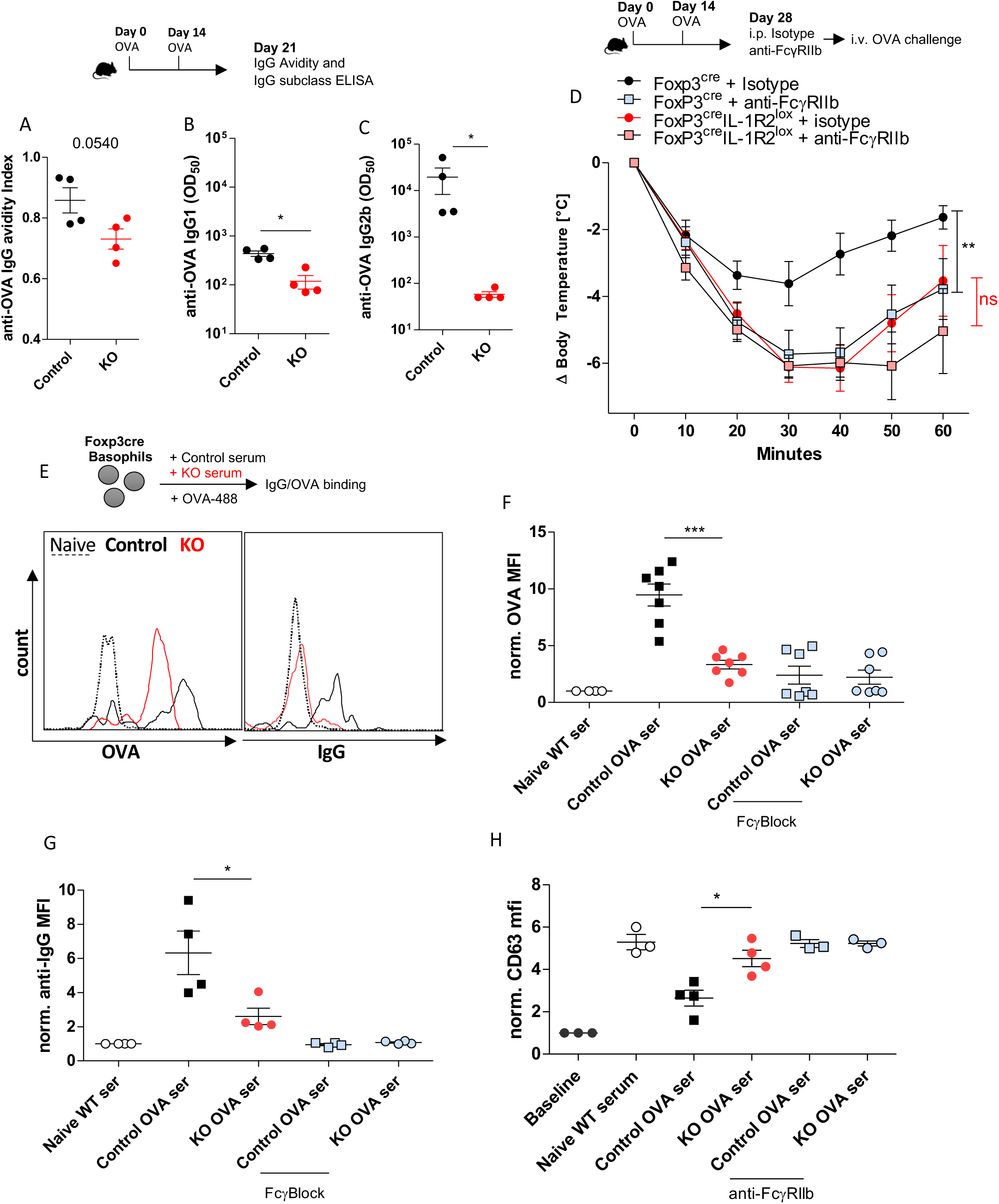
FoxP3^cre^IL-1R2^lox^ mice fail to suppress systemic anaphylaxis and basophil degranulation via FcγRIIb. The IgG response in FoxP3^cre^ (Control) or FoxP3^cre^IL-1R2^lox^ (KO) was assessed by ELISA. A) Shown is the mean mean ± SEM anti-OVA IgG avidity. B+C) OD_50_ titers of OVA-specific IgG1 (B), IgG2b (C). D) Mice were sensitized with OVA. 1 hour before OVA challenge, anti-FcγRIIb blocking antibody or isotype control antibody was injected. Shown is the mean ± SEM degrees lost compared to baseline temperature. E) Basophil assays were performed in the presence of serum. Shown are representative histograms for OVA-A488 or anti-IgG fluorescence to measure OVA-IgG immune complex binding. F, G) Shown are summarized mean ± SEM OVA-A488 MFI (F) or anti-IgG MFI (G). H) Basophil activation was inhibited with the presence of serum and FcγRIIb was blocked. Shown is the anti-CD63 MFI.

The IgG Fc Receptor FcγRIIb provides direct inhibitory signals for the IgE-dependent activation of allergic effector cells. To investigate the role of FcγRIIb in FoxP3^cre^IL-1R2^lox^ and their FoxP3^cre^ control mice *in vivo*, we blocked FcγRIIb prior to OVA challenge. As shown in Figure 4D, FcγRIIb blockade significantly increased anaphylaxis in control FoxP3^cre^ mice but not in FoxP3^cre^IL-1R2^lox^ mice, suggesting that in those mice, IgG fails to exert a protective effect on allergen challenge by engaging FcγRIIb.

To study the IgG functionality further, we studied the impact of serum IgG on IgG-OVA binding to FoxP3^cre^ control basophils. Fluorescently labelled OVA (OVA-488) was incubated with diluted serum at 4°C and binding was assessed by flow cytometry (titrations, see suppl. Figure 2). IgG binding was investigated by using an anti-IgG staining antibody. Figure 4E shows representative flow cytometry histograms.

We observed higher level of IgG-OVA immune complex (IC) binding to basophils in presence of FoxP3^cre^serum compared to FoxP3^cre^IL-1R2^lox^ serum, an effect that was inhibited by Fcγ blockade (Figure 4F, G). Thus, IgG-dependent IgG-OVA IC binding to basophils is reduced in serum derived from FoxP3^cre^IL-1R2^lox^. The residual Fcγ-independent OVA binding, that is higher in sensitized basophils than in naïve basophils, likely occurs through surface-displayed IgE (suppl. Figure 2).

To study the impact of serum IgG on basophil degranulation, we established a serum inhibition assay using FoxP3^cre^ control basophils, based on anti-CD63 measurement by flow cytometry (titrations, see suppl. Figure 2). As shown in Figure 4H, serum from FoxP3^cre^ mice inhibited basophil degranulation more than serum derived from FoxP3^cre^IL-1R2^lox^ mice. This inhibition of basophil degranulation could be fully reversed by FcγRIIb blockade.

In summary, allergic sensitization in FoxP3^cre^IL-1R2^lox^ mice is not only driven by increased IgE, but at the same time by a lack of protective allergen-specific IgG as in contrast to FoxP3^cre^ controls, FoxP3^cre^IL-1R2^lox^ mice fail to inhibit allergy through FcγRIIb engagement. In turn, IL-1R2 expression by Tfr may ensure the generation of IgGs that protect from allergy.

### FoxP3^cre^IL-1R2^lox^ mice display dysregulated GC B cell proliferation

Having shown the mechanisms that drive effector cell activation and anaphylaxis in FoxP3^cre^IL-1R2^lox^ mice, we next aimed to study the GC B cell response in further detail. We had previously observed that naïve Tfr/Tfh ratios are equal in FoxP3^cre^IL-1R2^lox^ whereas naïve GC B levels are increased^30^. To further investigate GC B levels upon sensitization, we isolated spleens from sensitized FoxP3^cre^ and FoxP3^cre^IL-1R2^lox^ mice and investigated their profile by flow cytometry. Gating strategies are displayed in suppl. Figure 3 or were previously shown^30,31^.

The frequency of Tfh and Tfr cells in CD4+ T cells were overall very similar. However, in alternative flow cytometry setups, we noticed that absolute cell numbers, including T cells, Tfh/Tfr, and B cells were higher across the board in FoxP3^cre^IL-1R2^lox^ mice (suppl. Figure 3). However, this general increase in T and B cells was not proportionate, as FoxP3^cre^IL-1R2^lox^ mice displayed significantly lower frequencies of splenic B cells compared to control mice, suggesting defects in the B cell compartment (Figure 5A).

**Figure 5:**
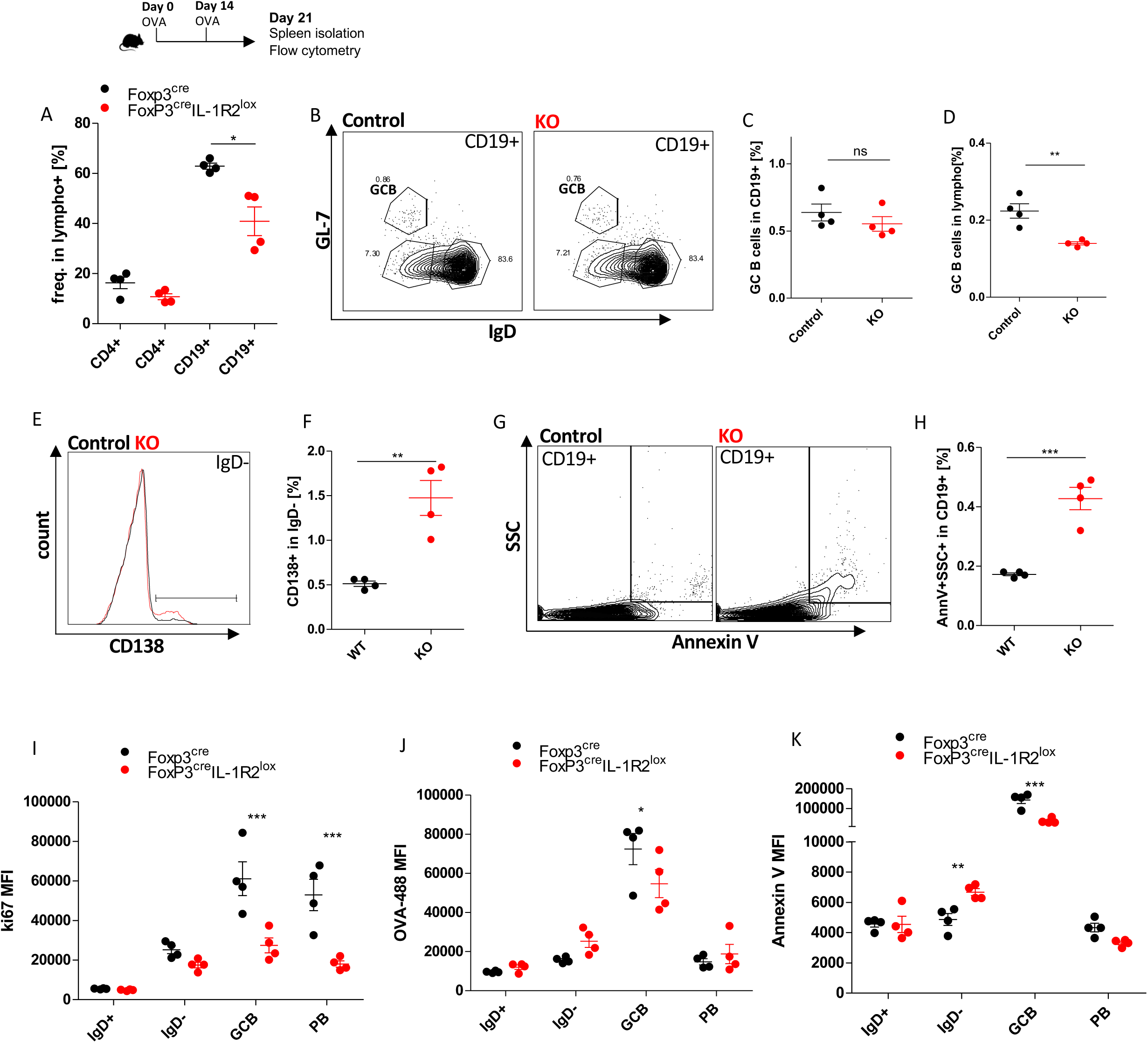
FoxP3^cre^IL-1R2^lox^ mice display dysregulated germinal center B cell proliferation. Foxp3^cre^ (Control) or Foxp3^cre^IL-1R2^lox^ (KO) mice were sensitized with OVA/Alum at day 0 and day 14. A) Mean ± SEM % of CD19+ and CD4+ in lymphocytes. B) Representative gating strategy for GCB cells. C) Mean ± SEM % of GCB cells in CD19+. D) Mean ± SEM % of GCB cells in lymphocytes. E) Gating strategy for plasmablasts F) Mean ± SEM % of plasmablasts in CD19+. G) Gating strategy for apoptotic B cells, H) Mean ± SEM % of apoptotic B cells. I-K) For IgD+ B cells, IgD-B cells, GCB and plasmablast (PB) subsets, shown are mean ± SEM MFI values of anti-ki67 staining (I), anti-AnnexinV staining (J), and OVA-MFI (K).

We next gated for GC B cells as shown in Figure 5B. Figure 5C shows that GC B cells did not change in frequency among FoxP3^cre^IL-1R2^lox^ mice but given the overall reduction of CD19+ B cells, GC B cell frequency was reduced in lymphocytes (Figure 5D).

In contrast to GC B cells, FoxP3^cre^IL-1R2^lox^ presented higher plasmablast frequencies as marked with CD138 staining (Figure 5E and 5F). This plasmablast increase likewise occurred in absolute numbers (suppl. Figure 3). When we investigated apoptotic B cells using Annexin V staining, we noted that frequency and absolute number of apoptotic B cells was increased in FoxP3^cre^IL-1R2^lox^ (Figure 5G and 5H, suppl. Figure 3).

Extensive GC B cell proliferation and apoptosis cycles are essential features in the process of selective hypermutation^34^. In a next step, we thus characterized proliferation/apoptosis profile across B cell stages. Additionally, we investigated OVA-488 binding to B cells in these subsets. Representative histograms for ki-67 staining (to visualize proliferation), as well as OVA-488/Annexin V staining are shown in suppl. Figure 3.

As shown in Figure 5I, the expression of proliferation marker ki-67 was reduced in FoxP3^cre^IL-1R2^lox^ B cell populations, most strikingly in GC B cells, but also in plasmablasts. Along the same lines, we observed that OVA-488 binding to GC B cells was lower in FoxP3^cre^IL-1R2^lox^ (Figure 5J). In contrast, the level of apoptosis in FoxP3^cre^IL-1R2^lox^ was overall higher in all IgD-cells but significantly lower in GC B cells (Figure 5K).

These findings suggest that the transition of activated B cells into proliferating antigen-specific GC B cells is disturbed in FoxP3^cre^IL-1R2^lox^ mice. Hence, IL-1R2 expression by Tfr may prevent the disruption of GC B cell maturation, which could explain the lack of allergen-specific IgG generated upon allergen immunization. Interestingly, this disruption does not seem to disrupt the generation of IgE and thus favours allergic sensitization.

### Tfr lacking IL-1R2 expression proliferate in response to IL-1β or antigen re-stimulation

Having mainly studied the effects of IL-1R2 deletion in Tfr on the allergic antibody response and B cells, we next aimed to investigate the underlying mechanisms in further detail. We have previously noted that Tfr/Tfh increase upon immunization in FoxP3^cre^IL-1R2^lox^ mice, although a specific mechanism was not identified^30^. We thus here aimed to test if the deletion of IL-1R2 in Tfr drives their own IL-1R1-dependent proliferation.

To functionally investigate this in closer detail, we established an *in vitro* re-stimulation experiment with splenocytes. The cells were cultured for 48 hours in presence of OVA and/or IL-1β. Strikingly, compared to splenocytes derived from control mice, follicular T cells in FoxP3^cre^IL-1R2^lox^ displayed a significant proliferative response to IL-1β, OVA and OVA+IL-1β re-stimulation (suppl. Figure 4).

Interestingly, the proliferation occurred only in Tfr, but not in Tfh cells (Figure 6A-C), which favours an increase of Tfr:Tfh ratios (suppl. Figure 4). We did not observe a proliferative response in Treg upon re-stimulation, in neither FoxP3^cre^ controls nor in FoxP3^cre^IL-1R2^lox^, which is in line with the previous findings that these subsets do not express IL-1 receptors. Hence, our findings suggest that deletion of IL-1R2 in FoxP3+ cells selectively enable IL-1 dependent proliferation in Tfr cells.

**Figure 6:**
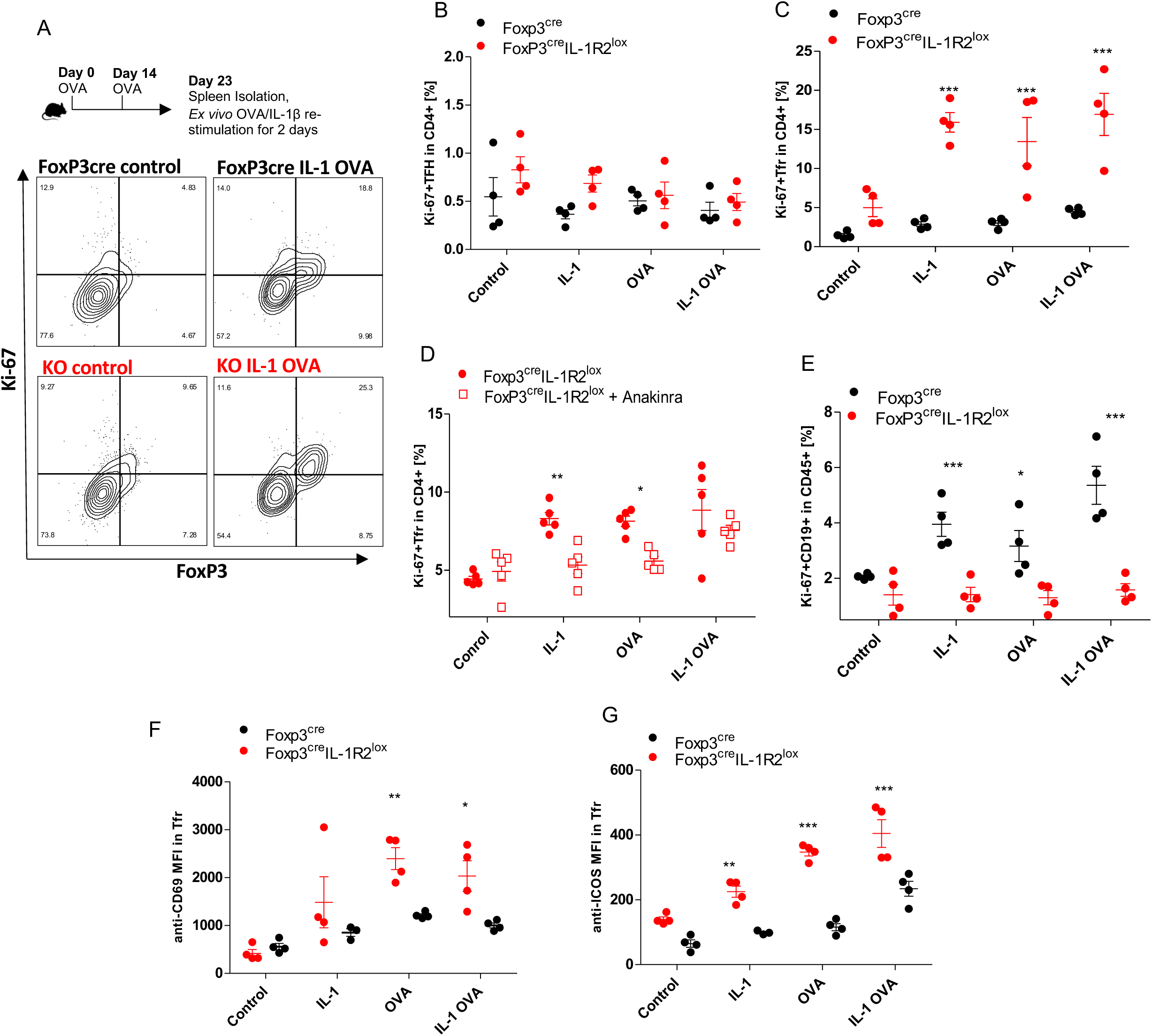
Tfr lacking IL-1R2 expression proliferate in response to IL-1β or antigen re-stimulation. At day 23, spleens were isolated and re-stimulated *ex vivo* with IL-1β, OVA or OVA+IL-1β for 48 hours. A) Shown is the gating strategy of proliferating Tfh/Tfr cells in flow cytometry. B) Mean ± SEM % of proliferating Tfh in CD4+. C) Mean ± SEM % of proliferating Tfr cells in CD4+. D) Anakinra was added to the 48-hour re-stimulation with IL-1β, OVA or OVA+IL-1β. Shown is the mean ± SEM % of proliferating Tfr cells in CD4+. E) Mean ± SEM % of proliferating CD19+ cells in CD45+ F) Shown is the mean ± SEM anti-CD69 MFI in Tfr. G) Shown is the mean ± SEM anti-ICOS MFI in Tfr.

To confirm the IL-1R1-dependency of Tfr proliferation, we performed an assay in the absence or presence of IL-1R1 antagonist Anakinra. As shown in Figure 6D, Tfr proliferation was blocked by the addition of Anakinra. When looking at the *in vitro* proliferation of B cells in our re-stimulation system, we did observe B cell proliferation in FoxP3^cre^ mice control that was completely suppressed in FoxP3^cre^IL-1R2^lox^ (Figure 6E). Hence, in line with *in vivo* results, *in vitro* B cell proliferation upon IL-1β and/or OVA re-stimulation was impaired in the presence of proliferating Tfr cells. When screening for Tfr-expressed molecules that change upon re-stimulation besides ki-67, we found and increase in their expression of CD69 and ICOS surface activation markers (Figure 6F, G).

In summary, *in vitro* re-stimulation of FoxP3^cre^IL-1R2^lox^ splenocytes selectively expands Tfr due to their ability to proliferate in IL-1R1 dependent fashion. These findings explain how Tfr/Tfh ratios are increased, and GC B cell responses are dysregulated in FoxP3^cre^IL-1R2^lox^ mice.

## DISCUSSION

We have previously highlighted the importance of Tfh/Tfr-expressed IL-1 receptors in the humoral immune response^29–31^. Our previous work concluded that Tfr are distinguished from Tregs by their expression of IL-1 receptors as opposed to IL-2 receptors^29–31^. However, the role of this axis in regulating antibody-driven pathologies such as allergy was not understood.

Here, we provide evidence that the IL-1 axis in GCs is an important regulator of the allergic response, unveiling the importance of IL-1R2 expression in Tfr cells. We demonstrate that targeted IL-1R2 deletion in FoxP3+ cells, which is expected to mainly target Tfr cells, drives allergic sensitization and anaphylaxis. Thus, IL-1R2 expression prevents the IL-1 dependent Tfr activation and thus prevents allergy exacerbation. Mechanistically, the allergic phenotype was driven by an increase of total IgE responses and by a quantitative and qualitative suppression of IgG responses, resulting in a lack of FcγRIIb engagement on allergic effector cells. This dysregulated antibody response may be derived from an increased IL-1/IL-1R1-dependent activation of Tfr, resulting in their own proliferation, and a suppression of GC B cell proliferation.

We have previously shown that the IL-1R1 is functional on Tfh cells, and indeed CD4^cre^IL-1R1^lox^ mice produced slightly lower levels of OVA-specific IgG than wild type controls upon OVA immunization^30,31^. At the same time IgG/IgE ratios were not significantly changed, and no difference in allergic phenotype was seen. These findings suggest that the allergic response can occur in absence of IL-1R1 expressing Tfh cells. A limitation of the model is that the effect of IL-1R1 deletion on Tfh is likely muted by expression of IL-1R2 in Tfr cells, suppressing IL-1-dependent Tfh activation by absorbing IL-1. Moreover, IL-1R1 can be expressed in both Tfh and Tfr subsets, although again, it seems likely that Tfr-expressed IL-1R2 would inhibit their own IL-1R1-dependent effects.

These effects are likely less of a problem for FoxP3^cre^IL-1R2^lox^ mice, because IL-1R2 expression is rather Tfr-specific, as neither conventional regulatory T cells (Tregs) nor Tfh express IL-1R2^29–31^. A recent study has found that Tfh can up-regulate FoxP3 in end-stage GC to shut down the reaction, thus essentially turning into Tfr cells^35^. A previous study has investigated IL-R2 on FoxP3-expressing cells and referred to them as Tregs^36^. However, most of these Tregs were found to display a “Tfr-like phenotype”, given their expression of PD-1 and CXCR5 whereas conventional IL-1R2+ Tregs were exceedingly rare. Nevertheless, despite all current evidence pointing towards the fact that Tregs do not express IL-1 receptors, we cannot exclude that this could be the case in certain circumstances. Here, the deletion of IL-1R2 expression in FoxP3+ cells activated IL-1R1-dependent Tfr but not Tfh or Treg proliferation, which is in line with our current hypothesis. Future studies will need to revisit the plasticity between these cell types and their phenotypes.

A recent study found that deletion of IL-1R2 enhances the overall GC response to immunization with SRBC due to an increase of IL-1 levels^37^. These findings are partly in line with ours, as we observe an overall increase of splenic cell number in FoxP3^cre^IL-1R2^lox^ mice. However, in our OVA/Alum sensitization model, the GC response and IgG levels were not enhanced but rather drastically suppressed, whereas only the IgE response was increased. Potentially, this important difference in GC B cell activation and IgE versus IgG response could be explained by the different antigen type/adjuvant system used for immunization.

The general importance of Tfr/Tfh in regulating allergic responses is increasingly recognized^22^. A recent study indicated that AIT modulates the Tfr/Tfh balance, highlighting a direct clinical relevance^38^. Overall, there have been conflicting reports on the mechanisms by which Tfr cells regulate the GC response in general and the allergic response specifically. An interesting study claimed that Tfr cells could promote allergy via IL-10^24^. We here could not confirm this link, but our study is in line with the overall narrative in that Tfr activation can promote allergy. We rather traced it to increased proliferative responses that may shift Tfr/Tfh and Tfr/B cell ratios. Given the variety of inhibitory tools that are per definition expressed by Tfr, it makes sense that their proliferation has a drastic effect on the GC reaction^39,40^.

We did observe an up-regulation of CD69, which has previously been shown to enhance suppressive function of FoxP3+ cells and its expression is regulated by NF-κB, a pathway that is triggered by IL-1 signaling^41,42^. The expression of CD69 was likewise observed in human Tfr located at the T-B border and was linked to Tfr suppressive activity^43^. The here-observed up-regulation of ICOS in IL-1 activated Tfr is in line with its previously described critical role in Tfr differentiation^44,45^. Future studies may shed more light on whether any individual Tfr-derived molecules are important in the regulation of allergic sensitization.

The induction and maintenance of the IgE responses is still a very debated topic. In contrast to specific IgG responses, total IgE levels do not necessarily require affinity maturation, but can occur rapidly and independently of GC^46^. Other studies argue that high-affinity GC responses are required for the induction of pathogenic IgE^46^. It is likewise still debated whether IgE class-switching occurs directly from IgM+ B cells or sequentially from IgG+ B cells^46^. A reason for these controversies is the fact that it is generally very difficult to IgE+ GC B cells^46^. Moreover, there are technical difficulties in distinguishing membrane IgE+ B cells from B cells that have bound IgE via the IgE receptor CD23^47^. In contrast to IgE+ GC B cells, IgE expression in immature plasmablasts is more common and associated with allergy^48–50^. Our findings can currently not solve these debates but we here show that in FoxP3^cre^IL-1R2^lox^, the GC response and affinity maturation is disrupted only for IgG, whereas the allergic IgE response can still occur. This dichotomy could be explained by a difference in GC kinetics for IgG versus IgE, or by that fact that IgE arises through a GC-independent mechanism. Future studies will focus more on unveiling those mechanisms.

While we know that human Tfr/Tfh express IL-1 receptors, our findings are currently limited to a mouse model of allergy. The study of circulating Tfr/Tfh is possible but has its own limitations as it does likely not reflect the situation within follicles. Another limitation that we could not address in this study is the role of IL-1Ra, which is likewise produced in Tfr cells^31^. The generation of Tfr-specific IL-1Ra knockout mice may delineate the role between IL-1Ra and IL-1R2 in Tfr cells in future studies.

The potential for clinical implications of our findings is obvious, as we are unravelling a novel role for IL-1R2-expressing Tfr in the prevention of allergic sensitization. Overall, the reasons why allergies develop are still insufficiently understood. However, we do know that natural and acquired allergen tolerance are often characterized by high IgG/IgE ratios^7^. Our findings suggest that IL-1R2 expression in Tfr could be an important factor that ensures the maintenance of high IgG/IgE ratios and thus prevents the onset of allergic sensitization. In contrast, a lack of IL-1R2 expression in Tfr during allergen encounter may favour allergic sensitization and exacerbate allergy. Down the line, the IL-1/IL-1R pathways in Tfr/Tfh during the GC response may be harnessed for novel diagnostic and/or therapeutic approaches in allergy.

In conclusion, we show that IL-1R2 expression in Tfr is an inhibitor of allergy as it limits the IL-1 dependent proliferative response of Tfr, which may disrupt GC B cell proliferation and suppress the IgG/IgE ratio.

## Notes

CONFLICT OF INTEREST: All authors declare no conflict of interest.

### Competing Interest Statement

The authors have declared no competing interest.

