## Supplementary material for "Expression of IL-1R2 by T follicular regulatory cells prevents the exacerbation of allergy by blocking their IL-1-dependent proliferation": Suppl. Figures

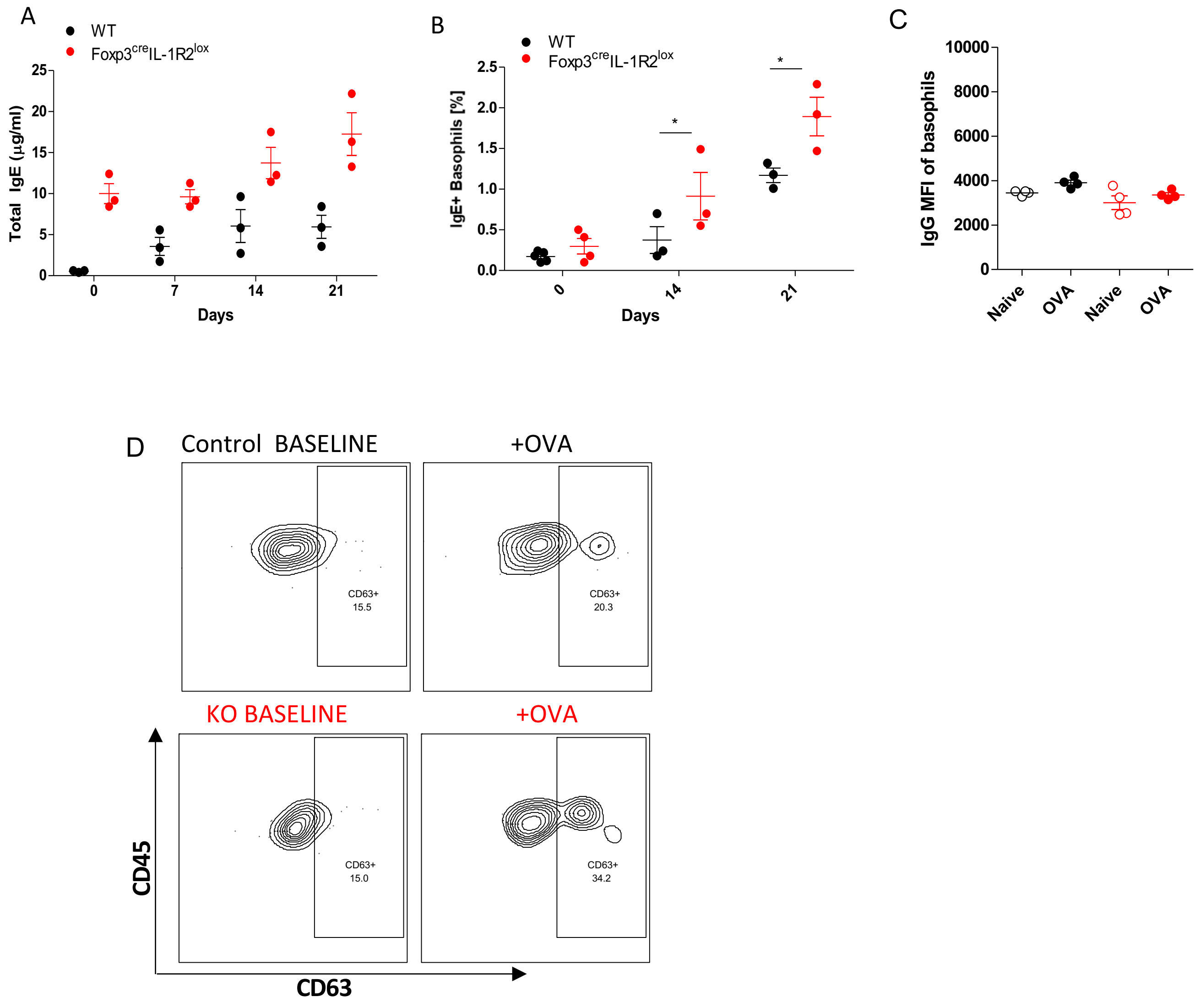


**Suppl. Figure 1:** A) Mice were sensitized with OVA/Alum. Total IgE levels in immunized FoxP3$^{cre}$ or Foxp3$^{cre}$IL-1R2$^{lox}$ mice over time. B) % IgE+ Basophils in immunized FoxP3$^{cre}$ or FoxP3$^{cre}$IL-1R2$^{lox}$ mice over time. C) Mean ± SEM basophil anti-IgG MFI at day 21. D) Second representative raw FACS plot showing basophil activation in FoxP3$^{cre}$ (Control) or FoxP3$^{cre}$IL-1R2$^{lox}$ (KO) mice.

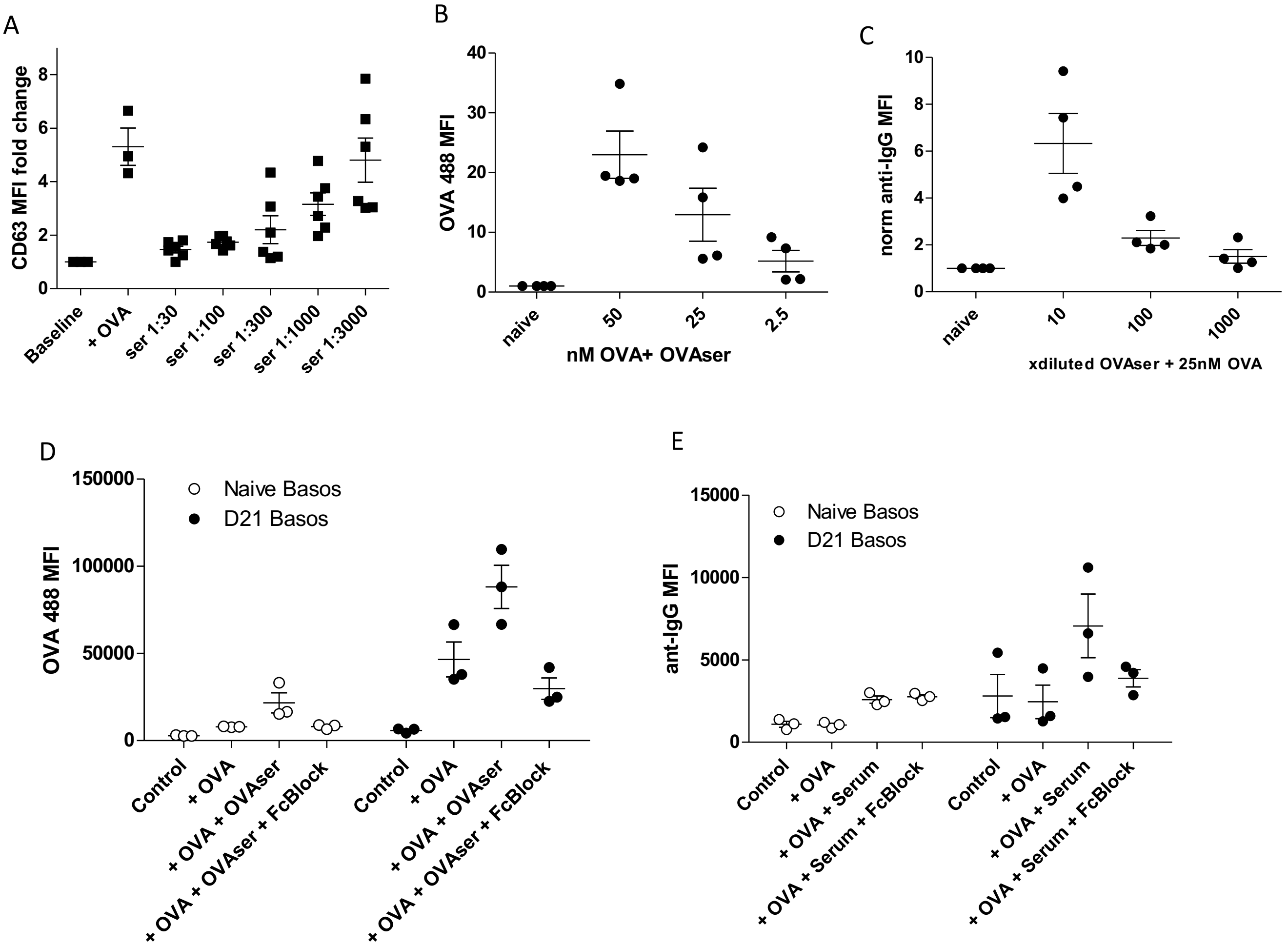


**Suppl. Figure 2:** Blood basophil assays were established. A) Mean ± SEM anti-CD63 MFI upon OVA with titrated serum from D21 OVA immunized mice. B) OVA-A488 titration to 1:10 serum from D21 OVA immunized mice. Shown is the mean ± SEM OVA-A488 fold change relative to naive serum. C) Serum from D21 OVA immunized mice titrated to 25nM OVA. Shown is the mean ± SEM anti-IgG MFI fold change relative to naive serum. D) OVA-A488 binding to naive basophils and D21 immunized basophils in additional presence of D21 OVA serum, and FcγBlock. Shown is the mean ± SEM OVA-A488 MFI. E) IgG binding to naive basophils and D21 immunized basophils inthe additional presence of D21 OVA serum, and FcγBlock. Shown is the mean ± SEM anti-IgG MFI.

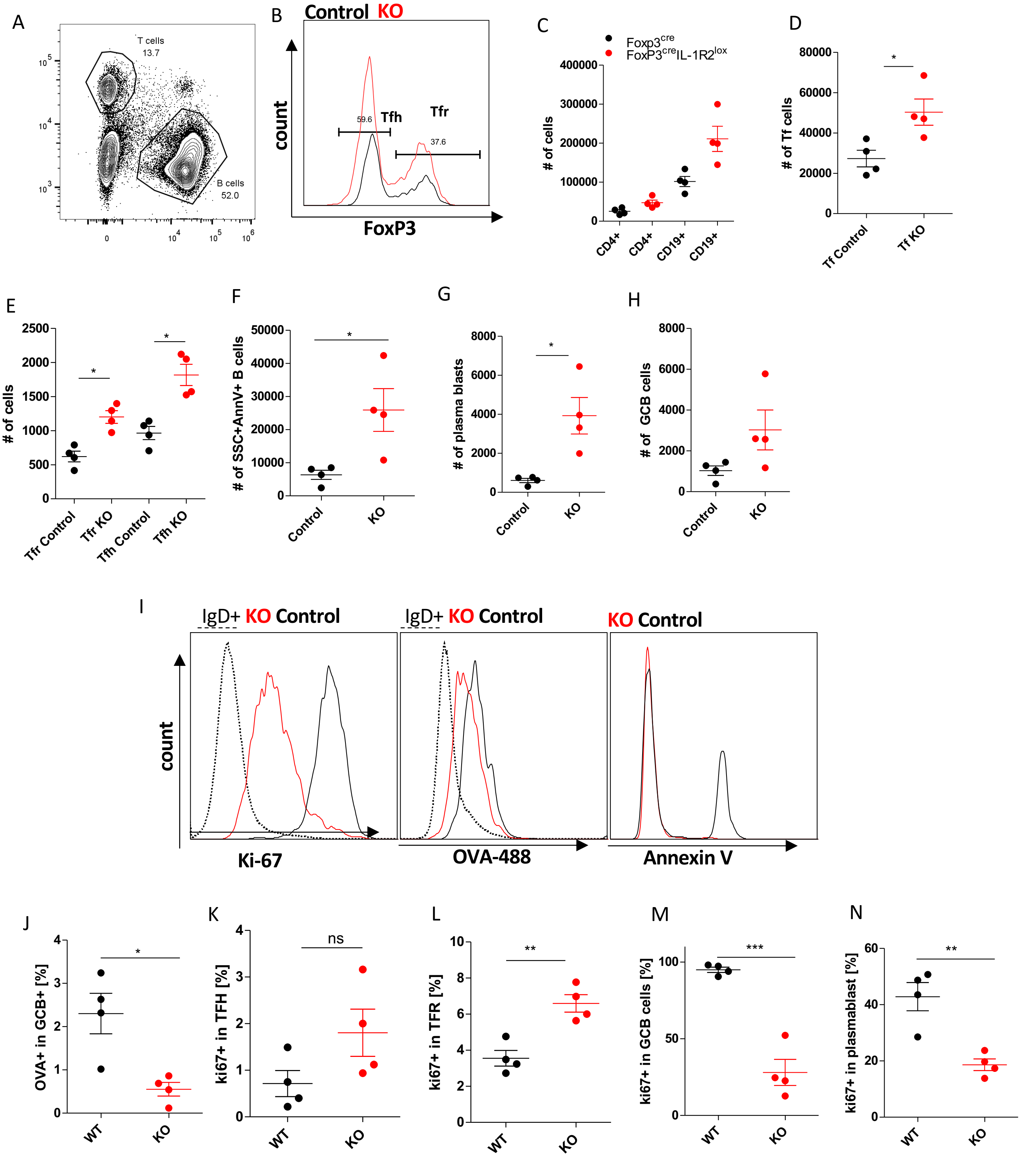


**Suppl. Figure 3**. WT or FoxP3$^{cre}$IL-1R2$^{lox}$ mice were sensitized with OVA/Alum at day 0 and day 14, spleens were investigated by flow cytometry. A) Gating strategy for CD4+ T cells and CD19+ B cells. B) Gating strategy for FoxP3+ cells in PD1+CXCR5+ cells. Shown are mean ± SEM, B cell/T cell numbers (C), Tf numbers (D), Tfh/Tfr numbers (E) apoptotic B cell numbers (F), plasmablast number (G), GC B cell numbers (H). Representative histograms for Annexin V, ki-67, and OVA-488 in GC B cells. J-N) Mean ± SEM % OVA+ in GC B cell (J), ki-67+ in Tfh (K), Tfr (L), GCB (M), and plasmablast (N).

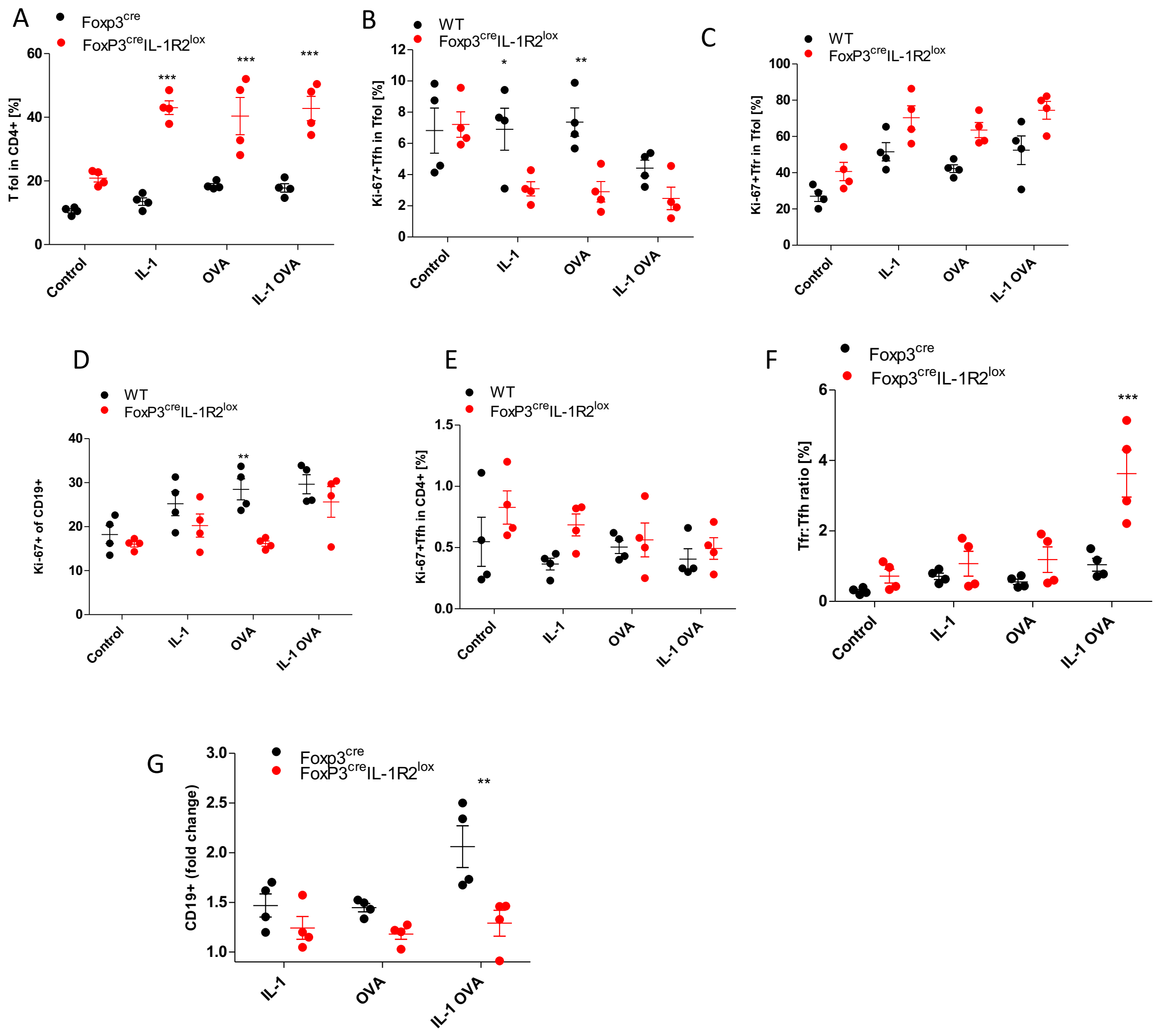


**Suppl. Figure 4**. FoxP3$^{cre}$ or FoxP3$^{cre}$IL-1R2$^{lox}$ mice were sensitized with OVA/Alum at day 0 and day 14. At day 23, spleens were isolated and re-stimulated *ex vivo* with IL-1β, OVA or OVA+IL-1β for 48 hours. A) Mean ± SEM % of proliferating Tfol cells in CD4+ B) Shown are mean ± SEM % of Ki-67+ Tfh in Tfol. C) Shown are mean ± SEM % ki-67+ Tfr in Tfol. D) Mean ± SEM % ki-67+ in CD19+. E) Shown are mean ± SEM % ki-67+ Tfh in CD4+ cells. F) Shown is the mean ± SEM Tfr/Tfh ratio. G) Shown is the mean ± SEM relative increase in B cell numbers.

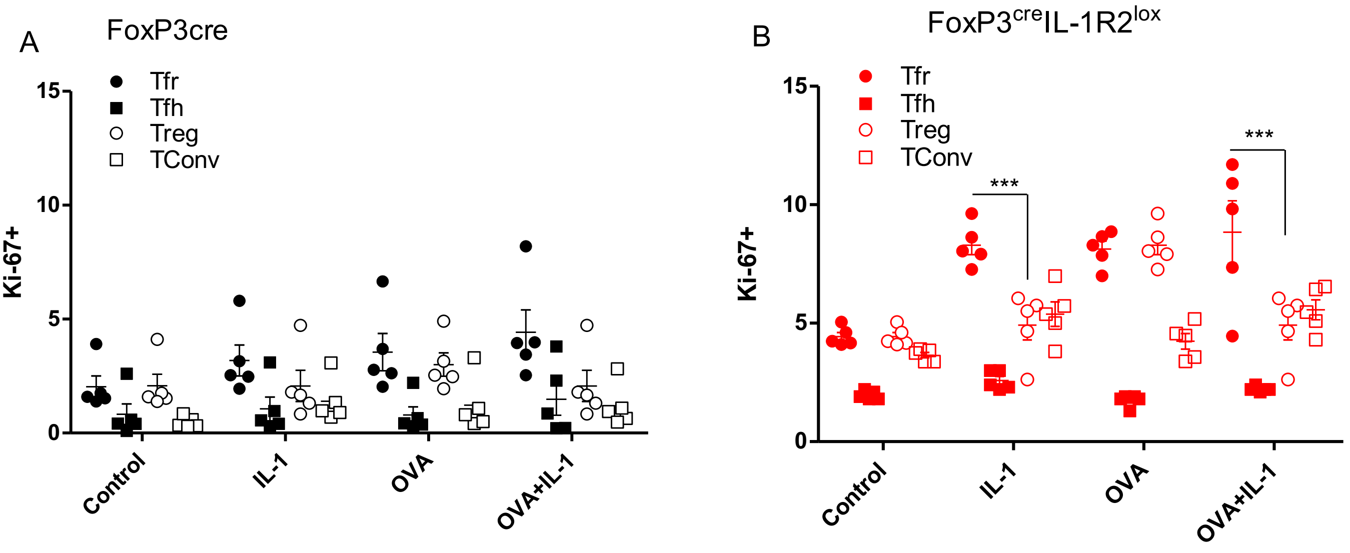


**Suppl. Figure 5**. FoxP3$^{cre}$ or FoxP3$^{cre}$IL-1R2$^{lox}$ mice were sensitized with OVA/Alum at day 0 and day 14. At day 23, spleens were isolated and re-stimulated ex vivo with IL-1β, OVA or OVA+IL-1β for 48 hours. A) Shown is the % of proliferating Tfr, Tfh, Treg and Tconv in response to re-stimulation in control FoxP3$^{cre}$ mice. B) Shown is the % of proliferating Tfr, Tfh, Treg and Tconv in response to re-stimulation in control FoxP3$^{cre}$IL-1R2$^{lox}$ mice.
